# Effects of substituents on anticancer activity of thiosemicarbazone against MCF-7 human breast cancer cell line

**DOI:** 10.1101/2020.02.19.955690

**Authors:** Belay Zeleke Sibuh, Pankaj Taneja, Sonia Khanna

## Abstract

**Background/Aim:** Breast cancer is one of the world’s leading cause of deaths in women. This study evaluated the in-vitro anticancer activity of different thiosemicarbazones (HacTSc, HSTsc, 3-MBTSc, 4-NBTSc and 4-HBTSc) against MCF-7 human breast cancer cell line and MCF-10 normal cell.

**Materials and Methods:** The ligands were prepared and characterized by UV vis, IR, ^1^H NMR. MTT assay was used to determine cell viability. Then data were analyzed using two-way ANOVA with Tukey post-hoc test. Result: The ligands have IC_50_ value ranging from 2.271µg/ml to 7.081µg/ml. Acetone thiosemicarbazone and 3-Methoxybenzaldehyde thiosemicarbazone, were identified as the most potent against MCF-7 breast cancer cells with IC_50_ value of 2.271µg/ml and 2.743µg/ml respectively. Whereas 4-Nitrobenzaldehyde thiosemicarbazone was the least potent. Also, the IC_50_ of the normal MCF-10 cell indicated their activity were selective. Conclusion: The activity of the ligands were dose, position and substituents dependent. Acetone thiosemicarbazone and 3-Methoxybenzaldehyde thiosemicarbazone are promising anticancer agents for further study.

## 1. Introduction

Thiosemicarbazones (hydrazine carbothioamides) are a family of compounds with high biological activity (1). Due to their versatile biological and pharmacological activities, their derivatives are of particular importance. They are good intermediates for pharmaceutical and bio-active material synthesis and are therefore, commonly used in medicinal chemistry field. They have also found their way into all branches of chemistry; they are used commercially as dyes, photographic films, plastic and textiles (2). In the past years, thiosemicarbazone derivatives have shown a wide range of biological activity *viz.* anti-fungal (3, 4), anti-tumor/cancer (5–10), Antimicrobial (11–13), sodium channel blocker (14), antiviral(15, 16).

Cancer is a malignant and invasive growth or tumor, especially of epithelium origin, which tends to recur after excision and may metastasize to other sites. It occurs due to failure of the mechanism that usually controls the growth and proliferation of cells (17). Cancer may affect people of all ages, and with increase in age the exposure to most type of cancer increases. Similarly, it can affect all parts of the body, for instance, lung and breast cancers are common all over the world (18).

Breast cancer is the most common type of cancer and the second leading cause of death among women worldwide (19). It affects 2.1 million women each year, and in 2018, an estimated 627,000 women died from breast cancer – about 15% of all female cancer deaths. While the rates of breast cancer among women are higher in developed region, rates are increasing globally in almost in every country (20).

Studies have shown that multi-drug resistance and unwanted side effects of current cancer chemotherapeutic contribute to an increased interest towards the development of new anticancer cancer drug, including synthetic compounds with minimal toxicity to normal tissue and highly effective with lower tumor cell resistance (21, 22). The present work is designed to evaluate the cytotoxic effects of different thiosemicarbazones on MCF-7 human breast cancer cell line.

## Material and Methods

All chemicals used were commercial products of analytical reagent grade procured from Sigma-Aldrich, except for the ligands, which were prepared by the reaction of MeOH and water solutions with different substituent and thiosemicarbazide using the procedures depicted below.

### Synthesis of Acetone Thiosemicarbazone (HacTsc)

Acetone thiosemicarbazone (C_4_H_9_N_3_S) was prepared by refluxing and stirring a mixture of acetone (0.02mmol, 1.16g) and thiosemicarbazide (0.02mmol, 1.822g) in 20ml distilled water. The mixture was refluxed for 15 min (40°c) and stirred for while resulting yellow colored solution, which was kept undisturbed. Then, a crystalline white product was separated out. Yield, 2.68g, Insoluble in: CHCl_3_, CCl_4_, H_2_O; soluble in DMSO, DHF/THF, CH_3_CN, ethanol, Methanol

### Synthesis of Salicylaldehyde Thiosemicarbazone (HSTsc)

Salicylaldehyde thiosemicarbazone, C_8_H_9_N_3_OS, was prepared by refluxing the mixture of salicylaldehyde (2.4 ml, 0.01mmol) and thiosemicarbazide (2.0g, 0.01 mmol) in a mixture of methanol (30 ml) and distilled water(30ml) as the solvent medium in the presence of glacial acetic acid (2-3 ml). The reaction mixture was refluxed for about 3-4 hours, resulting in a yellow colored solution, which was kept undisturbed. The next day, a crystalline yellow product was separated out. Yield, 4.046g, melting point 259°C. Insoluble in: CHCl_3_, CCl_4_, CH_3_CN, H_2_O; soluble in DMSO, DHF/THF

### Synthesis of 3-Methoxy benzaldehyde thiosemicarbazone (3-MBTsc)

3-Methoxy benzaldehyde thiosemicarbazone, C_9_H_11_N_3_OS, was prepared by refluxing a mixture of 3-Methoxy benzaldehyde (1.22 ml, 0.01mmol) and thiosemicarbazide (0.91g, 0.01 mmol) in a mixture of methanol (20ml) and distilled water(20ml) as the solvent medium in the presence of glacial acetic acid (2-3ml). The reaction mixture was refluxed for about 3-4 hours. Next day yellow crystalline compound separated out. Yield, 2.16g; Insoluble in: CHCl_3_, CCl_4_, CH_3_CN, H_2_O; ethanol, methanol; soluble in DMSO, DHF/THF

### Synthesis of 4-Nitrobenzaldehyde thiosemicarbazone (4-NBTSc)

4-Nitro benzaldehyde thiosemicarbazone was prepared by simple stirring 4-Nitrobenzaldehyde (1.51g, 0.01mmol) and thiosemicarbazide (0.914g, 0.01mmol) in a mixture of methanol (15ml) and distilled water (15ml) as a solvent medium. The reaction mixture was stirred for about 2 hours, resulting a yellow crystalline compound, which was kept undisturbed. Then a yellow crystalline compound was separate out. Yield, 2.30g; Insoluble in: CHCl_3_, CCl_4_, CH_3_CN, H_2_O, ethanol, Methanol; soluble in DMSO and DHF/THF

### Synthesis of 4-Hydroxybenzaldehyde thiosemicarbazone (4-HBTSc)

4-hydroxy benzaldehyde (4-HBTSc) was prepared by refluxing a mixture of 4-hydroxybenzaldehyde (1.222g, 0.01mmol) and thiosemicarbazide (0.914g, 0.01mmol) in a mixture of methanol (15ml) and distilled water (10ml) as a solvent medium. The reaction mixture was refluxed for about 5 hours, resulting a light-yellow colored solution, which was kept undisturbed. Then a light-yellow crystalline compound was separate out. Yield: 1.708g, melting point: 207°C; insoluble in CHCl_3_, CCl_4_, H_2_O, ethanol; soluble in DMSO, DHF/THF, CH_3_CN, Methanol

### Physical measurements

The structure of the ligand was confirmed from UV-Vis, infrared and ^1^H NMR spectral studies. UV Vis Spectroscopy was done by taking DMSO as reference sample or blank. The infrared spectroscopy was conducted to check whether the ligand is properly prepared or not. Also, ^1^H, nuclear magnetic resonance (NMR) spectra was measured in dimethyl sulfoxide (DMSO)-d_6_ as internal reference at ambient temperature. Also, the physical properties of the compounds were checked.

### Cell lines and cell culture

The test compound was studied for short-term *in vitro* anticancer activity against MCF-7 breast cancer cell line and for comparison, MCF-10 normal cell line was used. The cell lines were initially procured from the National Centre for Cell Science, Pune (NCCS). Cells were cultured in a 6 walled culture plate using Dulbecco’s Modified Eagle’s Medium, that was supplemented with 10% fetal bovine serum at 95% air condition with 5% of carbon dioxide (CO_2_) in 37°C Temperature.

### MTT cell viability assay Cell count

Before the cell viability test conducted cells were counted using hemocytometer (x10^4^cell/ml). Then cell viability was assessed by MTT assay using ATCC MTT Cell Proliferation Assay kit. MCF-7 and MCF-10 cell lines were plated separately in 96 well plates at a concentration of 1 × 10^4^ cells/well. After 48 hr. the media was removed and cells were washed twice with 100μl serum-free medium. Then 10μl MTT Reagent was added and left for incubation for 4 h at 37°C in a CO2 incubator, until intracellular purple formazan crystals are visible (until purple precipitate is visible). After that, MTT containing medium was discarded, the cells were washed with PBS (200μl). The crystals were then dissolved by adding 100μl of DMSO and mixed properly by pipetting up and down and left at room temperature in the dark for 2 hours. Spectro photometrical absorbance of the purple blue formazan dye was measured in a microplate reader at 570 nm. Then, percent Cell viability was calculated using the following formula.

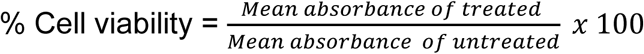

### Evaluation of anticancer potential

The in *vitro* anticancer activities of the test ligands were studied for short-period of time. First, 1mg of the synthesized thiosemicarbazone ligand was dissolved in 1 ml of Dimethyl sulfoxide (DMSO) separately. The test was done by taking the desired doses directly (without further dilution). Hence, the study was conducted in five concentrations including the control group (i.e. 0, 5µg/ml, 10 µg/ml, 12 µg/ml, 20 µg/ml of the test ligand) for each of the cell lines used. Based on the prepared test drug concentrations we have assigned the following groups. Group I: the cell line was subjected only for 10µl DMSO (control); Group II: 5µg/ml of the test ligand; Group III, 10µg/ml of the test ligand; Group IV, 12µg/ml test ligand; Group V, 20µg/ml test ligand.

### Statistical analysis

Data is presented in the form of descriptive statistics through tables and graphs. Descriptive Statistical analysis was done using Microsoft Excel. Whereas, IC_50_ value was calculated by fitting the sigmoidal dose response model (curve) by means of Prism statistical software package. Also, a two-way ANOVA and mean comparison test using Tukey multiple comparison test were analyzed in Prism statistical software package (GraphPad Software, Inc., La Jolla, CA, USA). *P*<0.05* was considered statistically significant.

## Results

The synthesis of different thiosemicarbazones were carried out according to the steps depicted in materials and methods section. The structures of the ligands were confirmed from UV-Vis, infrared and ^1^H NMR spectral studies.

### Anticancer activity of different thiosemicarbazone on MCF-7 breast cancer cell

MTT assay was used to test the anticancer activity of different substituent of thiosemicarbazone on MCF-7 breast cancer and MCF-10 normal cells. The half inhibitory concentration (IC_50_) of the test compounds was calculated using GraphPad prism software and presented in table1. According to the IC_50_ value, Acetone thiosemicarbazone by having IC_50_ value of 2.271 µg/ml is found to be the most potent and 4-Nitro benzaldehyde with IC_50_ value of 7.081 µg/ml is found to be the least potent. On the other hand, 3-methoxybenzaldehyde TSc (IC_50_=2.743 µg/ml) showed a comparable activity with Acetone thiosemicarbazone. On the other hand, the IC_50_ values of those ligands used in MCF-10 were higher than that of MCF-7 except in 4-nitro benzaldehyde where it is lower (2.255 µg/ml).

**Table 1.** IC_50_ values of the ligands against MCF-7 and MCF-10 cell lines

| Cell line | IC<sub>50</sub> µg/ml value |  |
| --- | --- | --- |
|  | MCF-7 | MCF-10 |
| Acetone TSc (HacTSc) | 2.271 | 115.8 |
| Salicylaldehyde TSc (HSTsc) | 3.361 | 5.399 |
| 3-methoxybenzaldehyde TSc (3-MBTSc) | 2.743 | 4.507 |
| 4-Nitrobenzaldehyde TSc (4-NBTSc) | 7.081 | 2.255 |
| 4-Hydroxybenzaldehyde TSc (4-HBTSc) | 3.605 | 9.55 |

### Acetone thiosemicarbazone inhibit the growth MCF-7 breast cancer cell line in lower concentration

The growth inhibition effect of **HacTSc** on MCF-7 and MCF-10 cells treated with different concentration was tested using MTT assay. The result (table 2 and figure 6A) suggested that, **HaTSc** inhibit approximately 6% percent of the tumor cell at concentration of 5 µg/ml and decrease to 3.5 % cell as the concentration increased to 20 µg/ml. The ligand’s toxicity to the normal cell showed cell viability increased as the concentration increase. This result suggests that Acetone thiosemicarbazone might be effective if it is used in very minimal concentration than high concentration.

**Table 2.**
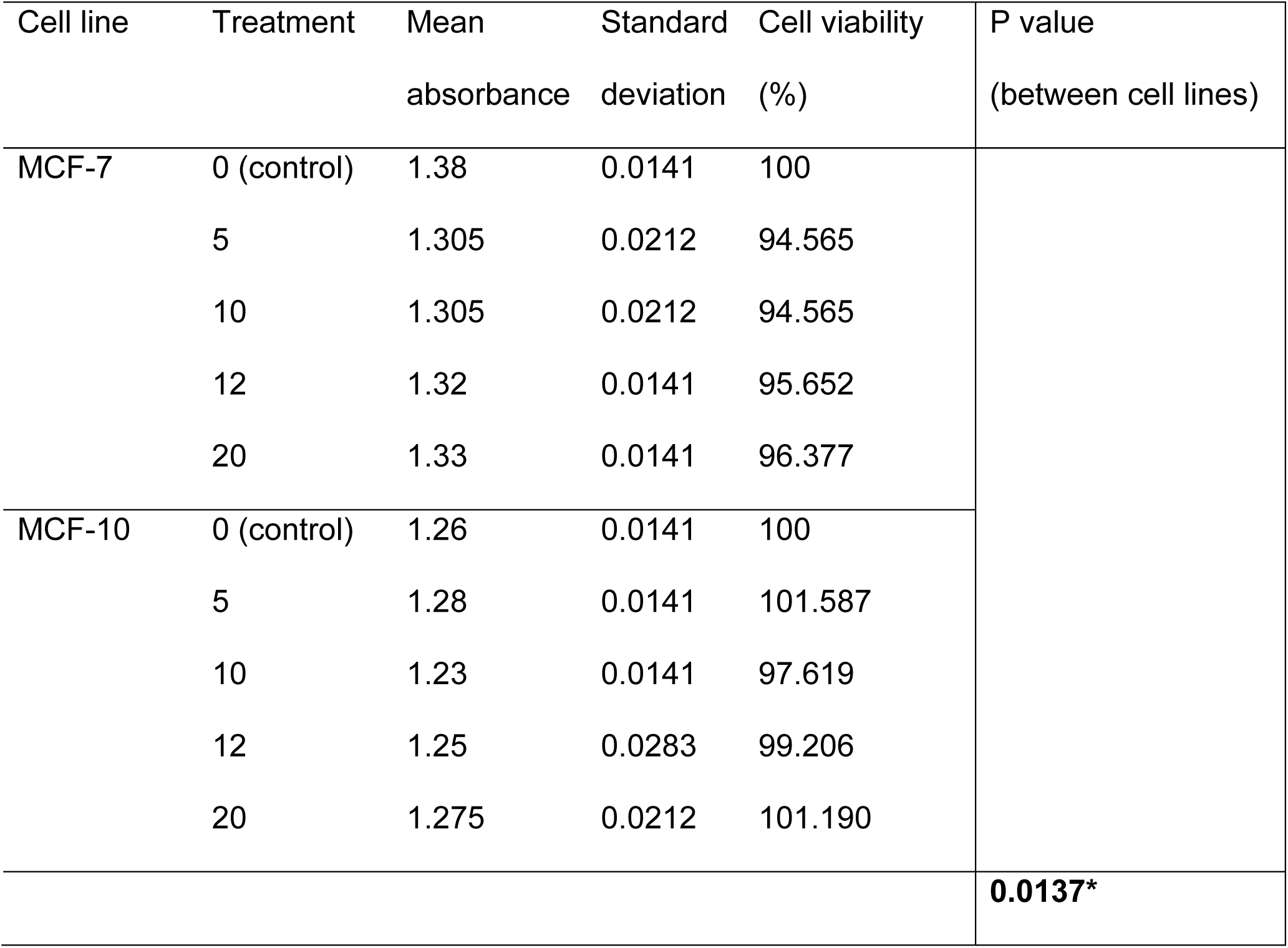
The effect of acetone thiosemicarbazone on cell viability

| Cell line | Treatment | Mean<br>absorbance | Standard<br>deviation | Cell viability<br>(%) | P value<br>(between cell lines) |
| --- | --- | --- | --- | --- | --- |
| MCF-7 | 0 (control) | 1.38 | 0.0141 | 100 |  |
|  | 5 | 1.305 | 0.0212 | 94.565 |  |
|  | 10 | 1.305 | 0.0212 | 94.565 |  |
|  | 12 | 1.32 | 0.0141 | 95.652 |  |
|  | 20 | 1.33 | 0.0141 | 96.377 |  |
| MCF-10 | 0 (control) | 1.26 | 0.0141 | 100 |  |
|  | 5 | 1.28 | 0.0141 | 101.587 |  |
|  | 10 | 1.23 | 0.0141 | 97.619 |  |
|  | 12 | 1.25 | 0.0283 | 99.206 |  |
|  | 20 | 1.275 | 0.0212 | 101.190 |  |
|  |  |  |  |  | **0.0137\*** |

**Figure 1.**
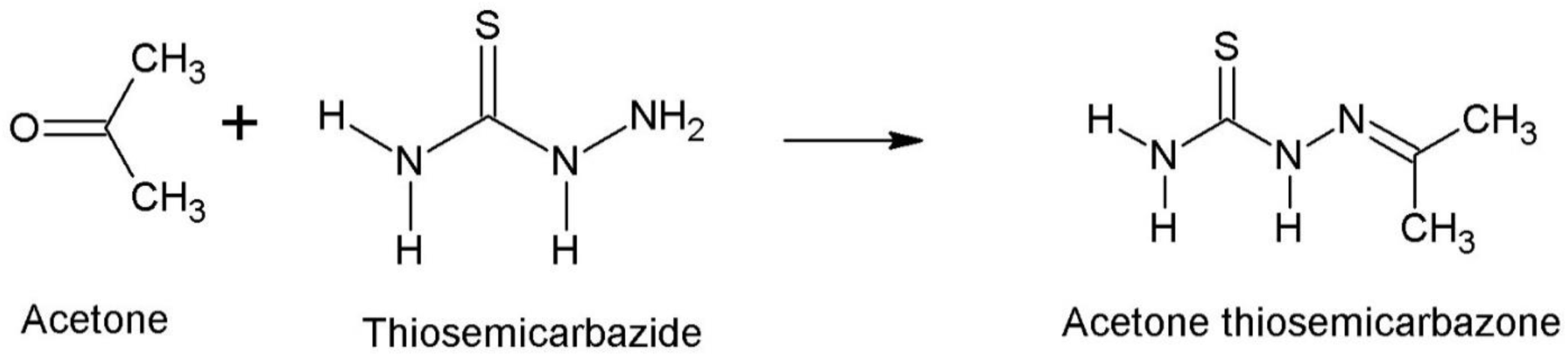
Preparation of Acetone thiosemicarbazone.

**Figure 2.**
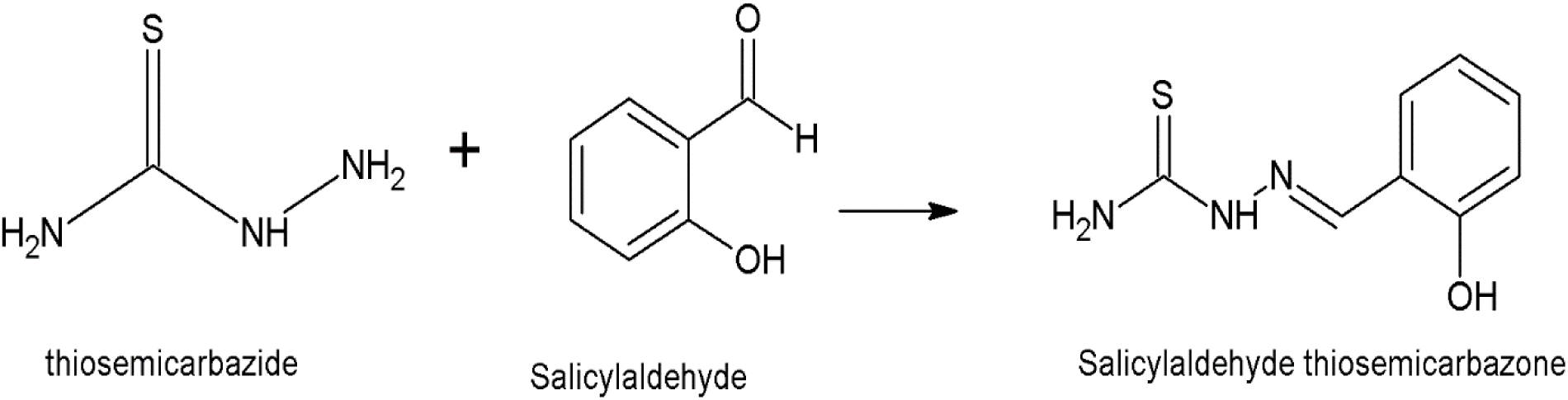
Preparation of salicylaldehyde thiosemicarbazone.

**Figure 3.**
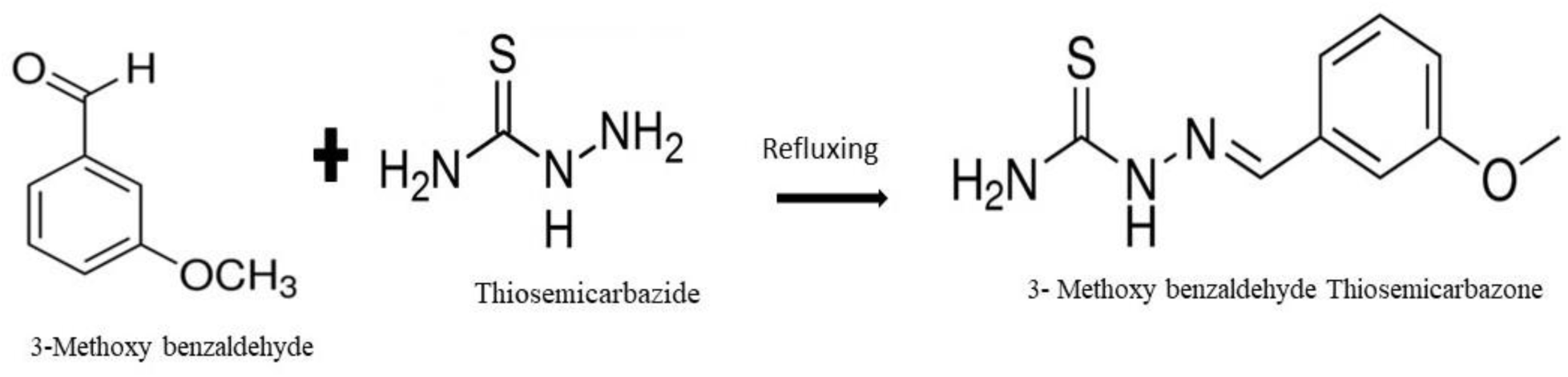
Synthesis of 3-Methoxy benzaldehyde thiosemicarbazone (3-MBTSc)

**Figure 4.**
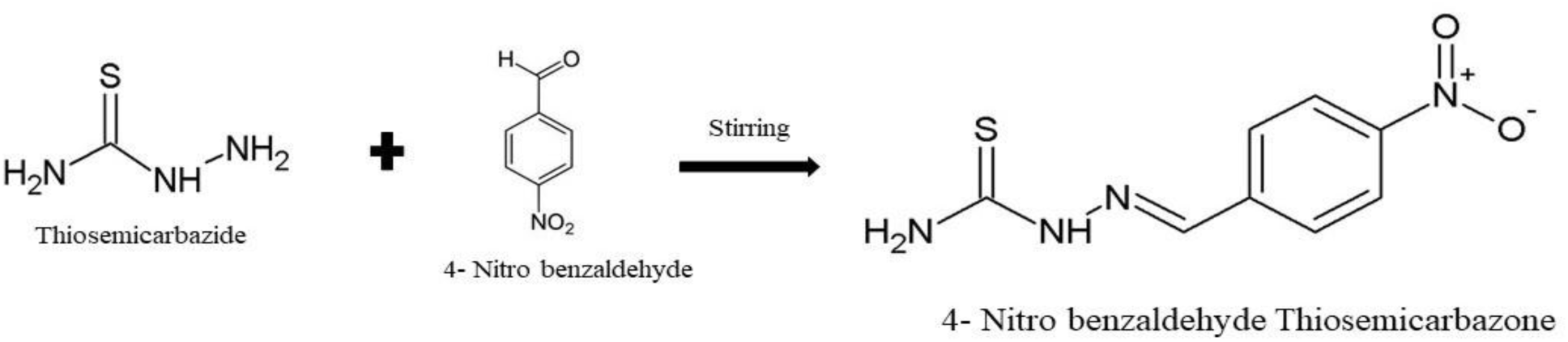
Synthesis of 4-Nitro benzaldehyde thiosemicarbazone.

**Figure 5.**
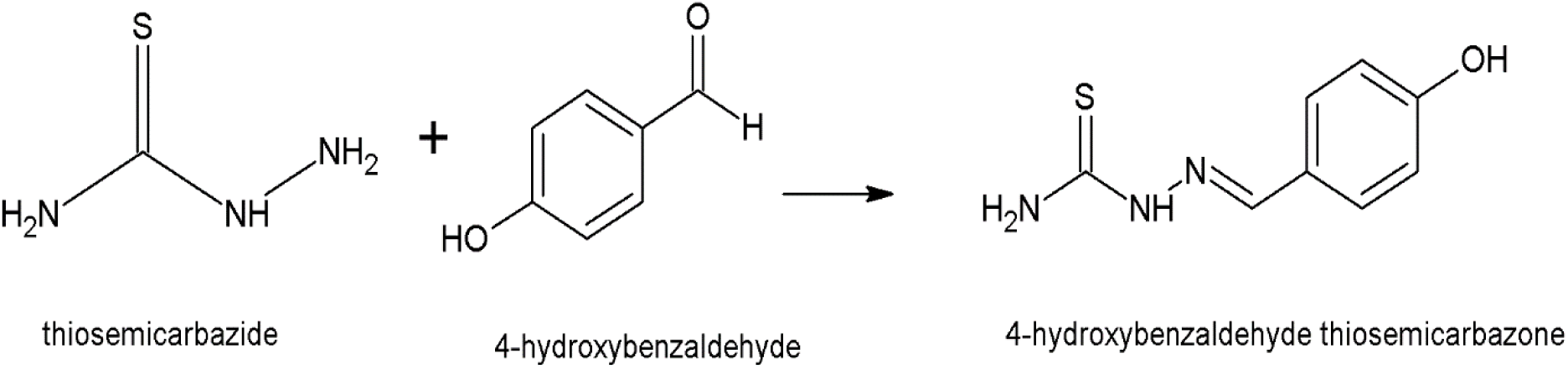
Preparation of 4-hydroxybenzaladehyde thiosemicarbazone.

**Figure 6.**
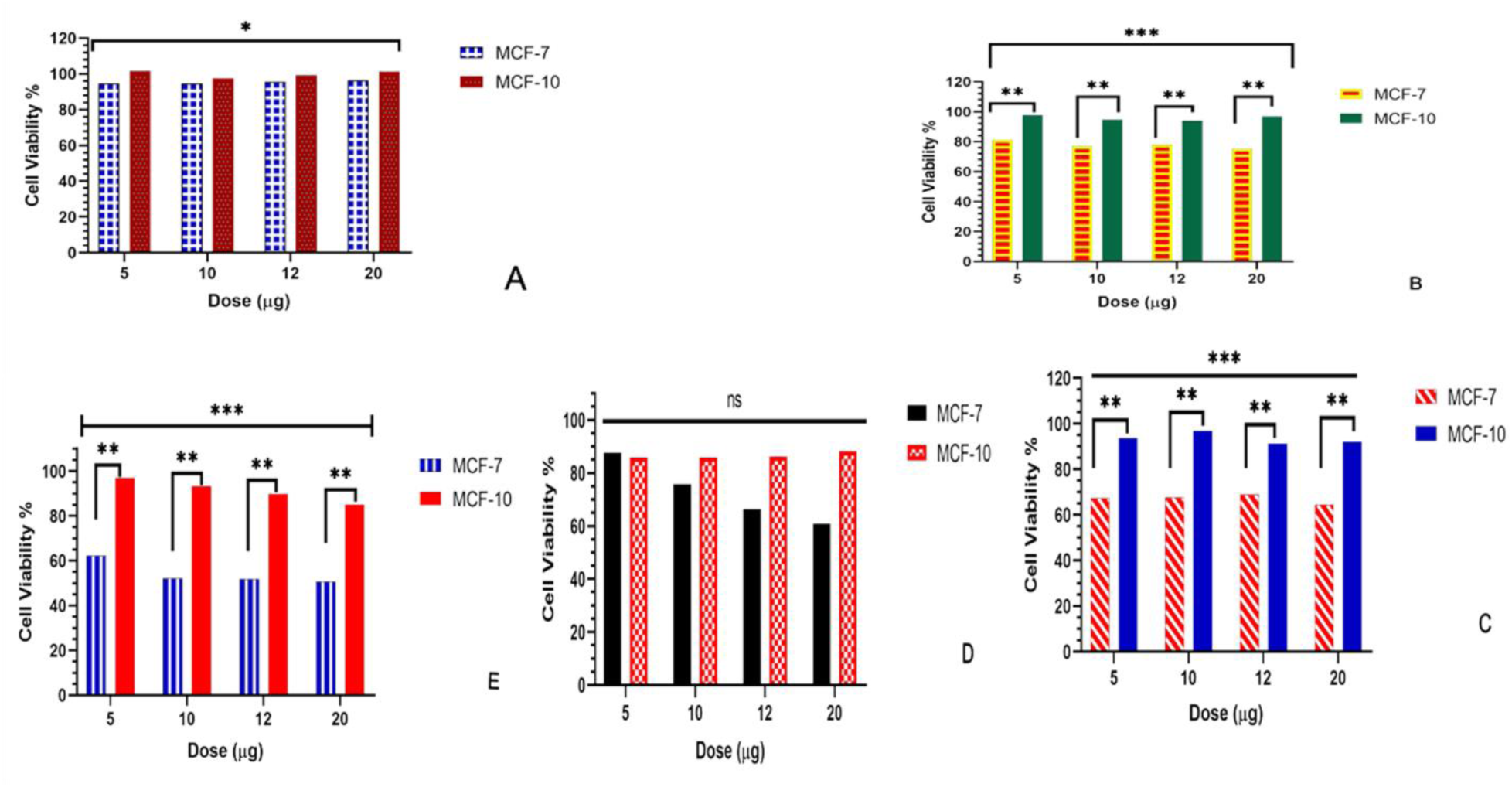
Effect of different thiosemicarbazone on cell viability (A) Acetone TSc (B) Salicylaldehyde TSc (C) 3-Methoxybenzaldehyde TSc (D) 4-Nitrobenzaldehyde TSC (E) 4-Hydroxybenzaldehyde TSC

### Salicylaldehyde thiosemicarbazone inhibit the growth MCF-7 breast cancer cell line

The MTT assay result showed that **HSTSc** has highly significant (p<0.001) growth inhibition on MFC-7 and MCF-10 cell. As depicted in table 3 and figure 6B, the test ligand showed 19% to 25% growth inhibition of MCF-7 when the concentration increased from 5 µg/ml to 20 µg/ml. On the other hand, MCF-10 growth inhibition suggest that the compound is less toxic to the normal cell with 97.6 % and 96.8% of viable cell after treatment of 5 µg/ml and 20 µg/ml respectively.

**Table 3.** The effect of Salicylaldehyde thiosemicarbazone on cell viability

| Cell line | Treatment | Mean<br>absorbance | Standard<br>deviation | Cell viability<br>(%) | P value<br>(between cell lines) |
| --- | --- | --- | --- | --- | --- |
| MCF-7 | 0 (control) | 1.38 | 0.0141 | 100 |  |
|  | 5 | 1.12 | 0.0141 | 81.159 |  |
|  | 10 | 1.06 | 0.0141 | 76.812 |  |
|  | 12 | 1.075 | 0.0212 | 77.899 |  |
|  | 20 | 1.04 | 0.0141 | 75.362 |  |
| MCF-10 | 0 (control) | 1.26 | 0.0141 | 100 |  |
|  | 5 | 1.23 | 0.0141 | 97.619 |  |
|  | 10 | 1.195 | 0.0071 | 94.841 |  |
|  | 12 | 1.185 | 0.0071 | 94.048 |  |
|  | 20 | 1.22 | 0.0283 | 96.825 |  |
|  |  |  |  |  | **0.0007\*\*\*** |

### Effect of 3-Methoxy benzaldehyde thiosemicarbazone on cell viability

As presented in table 4 and figure 6C, **3-MBTSc** showed a very promising and a significant (p<0.001) growth inhibition activity. It inhibits almost 33% of tumor cell growth at 5 µg/ml and continue up to 36.5% at 20 µg/ml. Comparing its toxicity to the normal cell at the same concentration the ligand inhibits 7-8% of MCF-10 cell growth.

**Table 4.** The effect of 3-Methoxy benzaldehyde thiosemicarbazone on cell viability

| Cell line | Treatment | Mean<br>absorbance | Standard<br>deviation | %cell<br>viability | P value<br>(between cell lines) |
| --- | --- | --- | --- | --- | --- |
| MCF-7 | 0 (control) | 1.38 | 0.0141 | 100 |  |
|  | 5 | 0.93 | 0.0141 | 67.391 |  |
|  | 10 | 0.935 | 0.0071 | 67.754 |  |
|  | 12 | 0.95 | 0.0141 | 68.841 |  |
|  | 20 | 0.89 | 0.0141 | 64.493 |  |
| MCF-10 | 0 (control) | 1.26 | 0.0141 | 100 |  |
|  | 5 | 1.18 | 0.0141 | 93.651 |  |
|  | 10 | 1.22 | 0.0141 | 96.825 |  |
|  | 12 | 1.15 | 0.0283 | 91.270 |  |
|  | 20 | 1.16 | 0.0141 | 92.063 |  |
|  |  |  |  |  | **0.0003\*\*\*** |

### The effect of 4-Nitro benzaldehyde thiosemicarbazone on cell viability

As demonstrated in table 5 and figure 6D, **4-NBTSc** inhibits around 13% of cells growth and continue up to 40% with increase in concentration from 5 µg/ml to 20 µg/ml. Also, the test ligand showed constant inhibition increment with small change in treatment concentration; at 10 µg/ml its effect was 25% and it increase by 9% and inhibit 34% of the cells while the dose is only 12 µg/ml. However, the ligand showed slight toxicity to normal cells and it inhibit 12-16 % MCF-10 cell growth. This output suggested that the activity of this ligand is promising but it might not be selective.

**Table 5.** The effect of 4-Nitro benzaldehyde thiosemicarbazone on cell viability

| Cell line | Treatment | Mean<br>absorbance | Standard<br>deviation | %cell<br>viability | P- value<br>(between cell lines) |
| --- | --- | --- | --- | --- | --- |
| MCF-7 | 0 (control) | 1.38 | 0.0141 | 100 |  |
|  | 5 | 1.21 | 0.0141 | 87.681 |  |
|  | 10 | 1.045 | 0.0071 | 75.725 |  |
|  | 12 | 0.915 | 0.0212 | 66.304 |  |
|  | 20 | 0.84 | 0.0141 | 60.870 |  |
| MCF-10 | 0 (control) | 1.26 | 0.0141 | 100 |  |
|  | 5 | 1.08 | 0.0141 | 85.714 |  |
|  | 10 | 1.08 | 0.0141 | 85.714 |  |
|  | 12 | 1.085 | 0.0071 | 86.111 |  |
|  | 20 | 1.11 | 0.0141 | 88.095 |  |
|  |  |  |  |  | **0.1176<sup>ns</sup>** |

### The effect of 4-hydroxy benzaldehyde thiosemicarbazone on cell viability

According to the MTT assay result (table 6 and figure 6E), **4-HBTSc** showed a very promising and highly significant (p<0.001) growth inhibition effect on MFC-7 and MCF-10 cells. It showed a 49.27% growth inhibition at dose of 20 µg/ml. Surprisingly at the concentration of 5 µg/ml it inhibits 38% of MCF-7 cells growth while it only inhibits 3% of MCF-10 normal cell. This result indicted that the compound is very promising for further study.

**Table 6.**
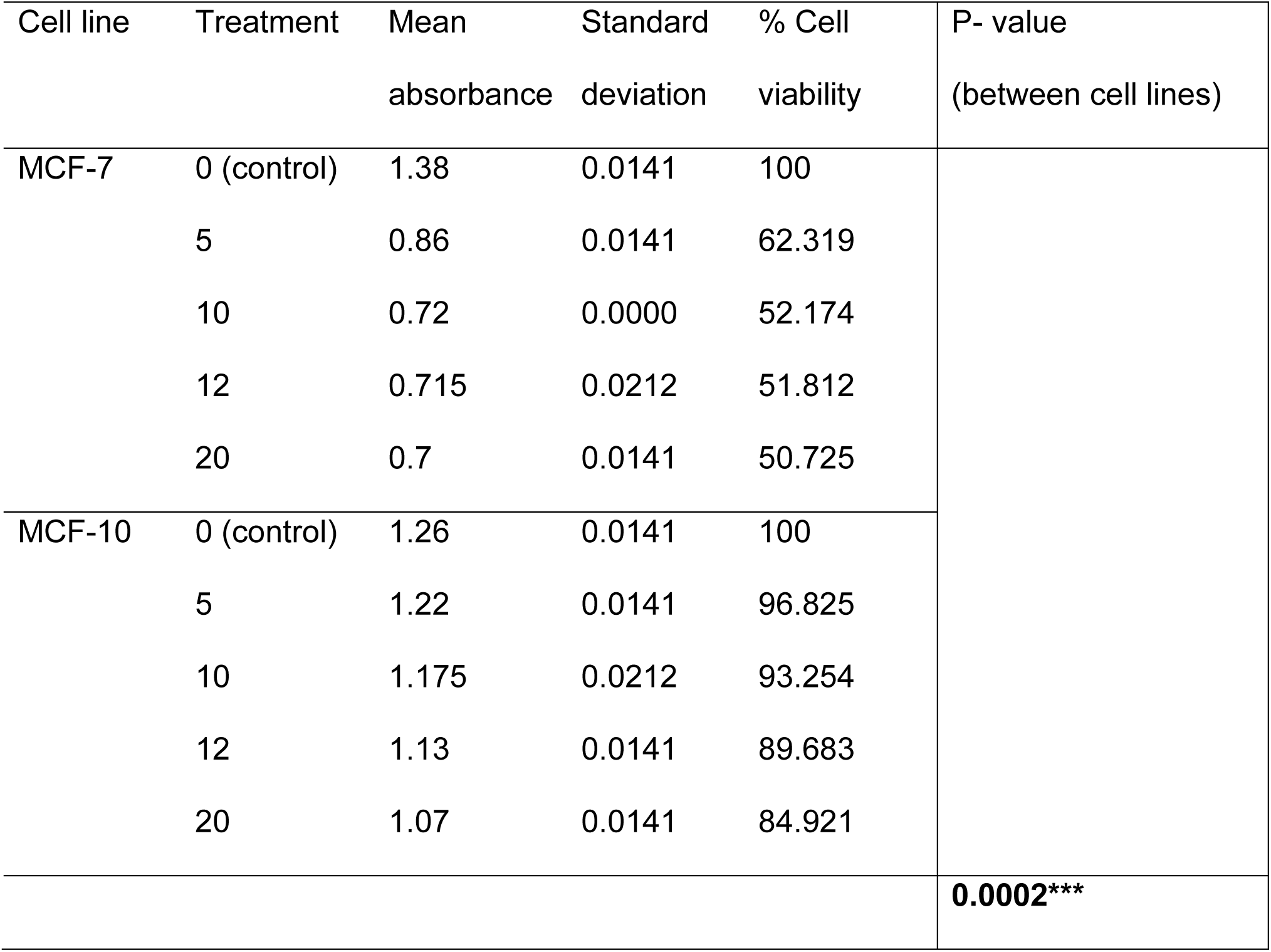
The effect of 4-hydroxy benzaldehyde thiosemicarbazone on cell viability

| Cell line | Treatment | Mean<br>absorbance | Standard<br>deviation | % Cell<br>viability | P- value<br>(between cell lines) |
| --- | --- | --- | --- | --- | --- |
| MCF-7 | 0 (control) | 1.38 | 0.0141 | 100 |  |
|  | 5 | 0.86 | 0.0141 | 62.319 |  |
|  | 10 | 0.72 | 0.0000 | 52.174 |  |
|  | 12 | 0.715 | 0.0212 | 51.812 |  |
|  | 20 | 0.7 | 0.0141 | 50.725 |  |
| MCF-10 | 0 (control) | 1.26 | 0.0141 | 100 |  |
|  | 5 | 1.22 | 0.0141 | 96.825 |  |
|  | 10 | 1.175 | 0.0212 | 93.254 |  |
|  | 12 | 1.13 | 0.0141 | 89.683 |  |
|  | 20 | 1.07 | 0.0141 | 84.921 |  |
|  |  |  |  |  | **0.0002\*\*\*** |

## Discussion

It was reported that structural variation, type substituent, cancer types and dose can affect the function of the compound (23–26). The IC_50_ and the ANOVA result in the current study also confirmed that, the activities of the ligands were significantly varied due to substituent type, dose and the type of cell line. As indicated in table 1, Acetone thiosemicarbazone with the IC_50_ value of 2.271 µg/ml is found to be the most potent and 4-Nitro benzaldehyde with IC_50_ value of 7.081 µg/ml is found to be the least potent. Acetone, which is the simplest and the smallest Ketone as a substituent showed a better anticancer activity than all aldehyde substituents used for this study. This could be correlated with the location of the substituent, as acetone is positioned with non-aromatic nature, while the other ligand substituents are located in the different position of the benzene ring. Conversely, among the benzene ring substituents we synthesized the methoxy group in 3-methoxy benzaldehyde thiosemicarbazone showed a better activity than the Nitro and hydroxy substituent. In fact, when we see the activity of the hydroxy group on its own, the effect of the substituent position is found to be lower.

Similarly, MCF-10 normal cell line was used to determine whether the test compounds were toxic or nontoxic to healthy cells. The IC_50_ values of those ligands used in MCF-10 were higher than that of MCF-7 except in 4-nitro benzaldehyde where it is lower. These results generally indicated that most of the ligands are selective in their anticancer effect. Particularly Acetone thiosemicarbazone showed highly selective activity with an IC_50_ value of 115.8 µg/ml against the normal cell line. On the contrary, 4-nitro benzaldehyde showed highly toxic activity with IC_50_ value of 2.255 µg/ml to the normal cell line.

The cell viability result revealed that the inhibition of cell growth significantly increased with an increase in dose and varies with a substituent except for Acetone thiosemicarbazone where the inhibition has decreased as the dose increases (Table 2 to 6). In MCF-7 breast cancer cell line, the highest percentage of inhibition (i.e. 49.27%, 48.19 and 47.83%) was observed on cells treated with 4-HBTSc at the dose of 20, 12 and 10 µg/ml respectively. The other ligands also showed a mild and comparable effect. In line with this, with an increase in dose, **4-NBTSc and 3-MBTSc** showed 40% and 35 % of growth inhibition respectively (table 4 and 5). On the contrary, the lowest growth inhibition effect was observed at doses of 20 µg/ml (table 2) due to **HacTSc**. Theoretically, the lowest IC_50_ is associated with the highest anticancer activity, and the highest cell growth inhibition. Thus, in the current study most of the percentage of cell death due to inhibition of cell growth is consistent with the IC_50_ value of the test ligands. Nevertheless, when we see the effect of Acetone thiosemicarbazone, its inhibition of growth decreased as the dose increased (5.4% to 3.6%). This could mean that **HacTSc** could be effective if it is used at small concentration. Generally, all test ligands showed significant (p<0.05*, P<0.001***) growth inhibition between MCF-7 and MCF-10 cells except **4-NBTSc** which showed statistically non-significant effect on cell viability between tumor and normal cells. This could be due to the toxic nature of **4-NBTSc** to the normal cell (MCF-10) which inhibits growth in close range with MCF-7 breast cancer cell. In conclusion, the test ligands showed potential as anti-tumor agents for further in-vitro and/or in-vivo stages of screening, encouraging further research in this field.

## Authors’ Contributions

All authors contribute equally in Research concept and design, data collection, Data analysis and interpretation, writing; critical revision and Final approval of the article.

## Acknowledgement

The authors would like to acknowledge Sharda university for their material support. Also, Belay Zeleke would like to thank Arba Minch University, Ethiopia, for sponsoring his Ph.D. study.

